# Cell-fate decision of mesenchymal stem cells toward osteocyte differentiation is committed by cell condensed condition in spheroid culture

**DOI:** 10.1101/2020.12.09.417097

**Authors:** Jeonghyun Kim, Taiji Adachi

**Affiliations:** Institute for Frontier Life and Medical Sciences, Kyoto University, Kyoto 606-8507, Japan

**Keywords:** Bone, osteocyte, spheroid, 3D culture, differentiation, mesenchymal stem cells

## Abstract

Osteocytes are mechanosensory commander cells to regulate bone remodeling throughout the lifespan. While the osteocyte is known as a terminally differentiated cell derived from mesenchymal stem cell, the detailed mechanisms of cell-fate decision toward osteocyte differentiation still remain unclear. In this study, we fabricated three-dimensional (3D) self-organized spheroids using human mesenchymal stem cells (hMSCs). Under the osteogenesis induction medium, the spheroid culture model exerted the osteocyte-likeness within 2 days compared to a conventional 2D monolayer model. By using an inhibitor of actin polymerization, we showed an involvement of actin balancing in the osteocyte differentiation in the spheroid. Notably, we represented that the cell condensed condition acquired in the 3D spheroid culture model determined a differentiation fate of MSCs to osteocytes via actin balancing. Taken together, we suggest that our self-organized spheroid model can be utilized as a new *in vitro* model to represent the osteocyte differentiation process and further to recapitulate an *in vitro* ossification process.

## Introduction

The bone undertakes continuous remodeling by osteoclasts and osteoblasts throughout the lifespan while its balance is modulated by mechanosensory commander cells, osteocytes (Robling and Bonewald, 2020; Adachi et al., 2009). From the view of developmental process, the ossification is initiated by mesenchymal condensation. In other words, the osteocyte is also known as a terminally differentiated bone cell derived from the mesenchymal stem cells (MSCs). The MSCs are referred to as multipluripotent progenitor cells capable of differentiation into skeletal tissues such as osteoblasts, chondrocytes, adipocytes, myocytes (Caplan, 1991; Pittenger et al., 2019, 1999). After the osteoblasts produce the bone matrices such as collagen type I and alkaline phosphatase during the bone remodeling process (Parfitt, 2001; Rutkovskiy et al., 2016), it is known that they encounter one of fates; 1) to become bone-lining cells, 2) to become osteocytes embedded in the bone, and 3) to undertake apoptosis (Manolagas, 2000; Robling and Bonewald, 2020).

For both osteoblast differentiation and bone mineralization studies, many researches have established a gold standard protocol to induce *in vitro* osteogenesis using chemical osteogenesis induction (OI) supplements such as ascorbic acid and β-glycerophosphate (Buttery et al., 2001; Coelho and Fernandes, 2000). On the other hand, *in vitro* osteocyte differentiation has been also attempted by many studies. After the osteoblast precursor cells were subjected to a long period of cultivation in the OI culture medium on the culture dish, a three-dimensional dome-shape of bone-like nodule was formed (Bhargava et al., 1988; Kawai et al., 2019; Mechiche Alami et al., 2016). Although the osteocyte-like cells were eventually observed inside the nodule after several weeks to months, the efficient method to achieve the direct osteocyte differentiation derived from pre-osteoblast cells or stem cells has not been established since the osteocyte-like cells were randomly shown in the nodule after such a long cultivation. Moreover, it still remains difficulties for primary osteocytes to culture *in vitro* and further to maintain the cellular morphology and functions for a long period of time, so that a new *in vitro* osteocyte model is required to be established with a novel efficient method. After achieving this, it will help unraveling the unknown mechanisms of cell-fate decision toward osteocytes.

In previous studies, our group has elucidated the effect of three-dimensional culture for pre-osteoblast cells on the osteocyte differentiation (Kim et al., 2020; Kim and Adachi, 2019). Compared to the conventional monolayer model, the pre-osteoblast cells in the form of scaffold-free spheroid model rendered the osteocyte-likeness within 2 days in the absence of the OI supplements. In this study, we fabricated the 3D spheroid culture model reconstructed by the MSCs and evaluated its osteocyte differentiation capability. By using the MSC spheroid model, we attempted to elucidate the mechanism of cell-fate decision of MSCs toward osteocytes and also to recapitulate the *in vitro* ossification process derived from the mesenchymal condensation.

## Materials and Methods

### Cell culture

Human bone marrow-derived MSCs (passage 2) were acquired from PromoCell (Germany) and RIKEN BRC (Japan). The cells were maintained in the basal medium which is low glucose-Dulbecco’s Modified Eagle Medium (Gibco) supplemented with 10% fetal bovine serum (Gibco) and 1% antibiotic-antimycotic (Gibco) solution in a humidified incubator at 37□ under 5% CO_2_ condition. We carried out the cell passaging every 2 – 3 days when the cell confluency became 80 – 90%. For experiments, the cells from passage 4 up to 12 were used. To prepare an osteogenic induction (OI) medium, we utilized high glucose-Dulbecco’s Modified Eagle Medium (Gibco) containing 50 μM ascorbic acid (Wako), 10 mM β-glycerophosphate (Sigma), and 100 nM dexamethasone (Nacalai Tesque). To prepare for 2D monolayer samples, 200,000 cells were subcultured on a 35 mm diameter culture dish to become confluent after 2-day incubation. This study was approved by the Ethics Committee of Institute for Frontier Life and Medical Sciences, Kyoto University (Clinical Protocol No. 89).

### Fabrication of spheroids

Based on the previous study (Kim et al., 2020), we fabricated self-organized spheroids using hMSCs on the Nunclon Sphera 96-well round-bottom plate (ThermoFisher). 2,500 cells were subcultured on the round-bottom plate to fabricate the spheroids and incubated for 2, 4 or 7 days in the presence or absence of the OI medium depending on the aims of each experiments.

### Real time-PCR

To measure mRNA expressions in the spheroid and monolayer reconstructed by hMSCs, we conducted real-time PCR. After the samples were rinsed with PBS once, they are immediately lysed with Isogen II (Nippon Gene). According to the manufacturer’s protocol, RNA extraction and cDNA synthesis were carried out using PureLink RNA Mini kit (Invitrogen) and Transcriptor Universal cDNA Master (Roche), respectively. For the real-time PCR, we used PowerUp SYBR Green Master Mix (ThermoFisher) while all the PCR primers were designed as shown in Table 1. *Nanog*, *Oct4*, and *Sox2* were used as undifferentiated stem cell markers. For osteoblast markers, we examined mRNA expressions of runt-related transcription factor 2 (*Runx2*), alkaline phosphatase (*Alpl*), and type I collagen, alpha 1 chain (*Col1a1*). To distinguish osteocytes, we utilized osteopontin (*Opn*), phosphate regulating endopeptidase homolog X-linked (*Phex*), and sclerostin (*Sost*). While SRY-box transcription factor 9 (*Sox9*) and aggrecan (*Acan*) were used as chondrocyte markers, type X collagen, alpha 1 chain (*Col10a1*) and matrix metallopeptidase 13 (*Mmp13*) were examined as hypertrophic chondrocyte markers. For adipocyte markers, we utilized peroxisome proliferator activated receptor gamma (*Pparg*) and CCAAT enhancer binding protein alpha (*Cebpa*). All the mRNA expressions were normalized to a reference gene, glyceraldehyde-phosphate dehydrogenase (*Gapdh*) and measured by using the 2^−ΔΔCT^ method.

**Table 1.**
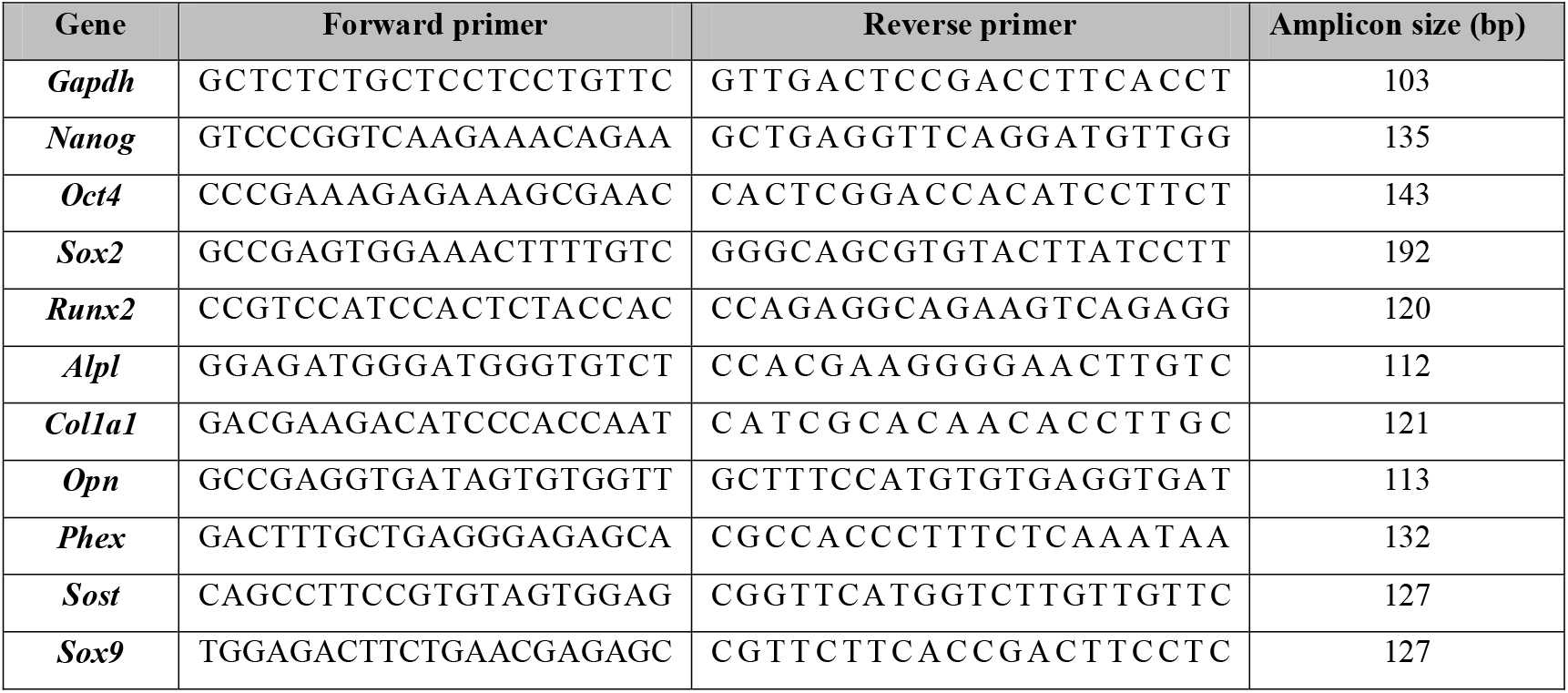
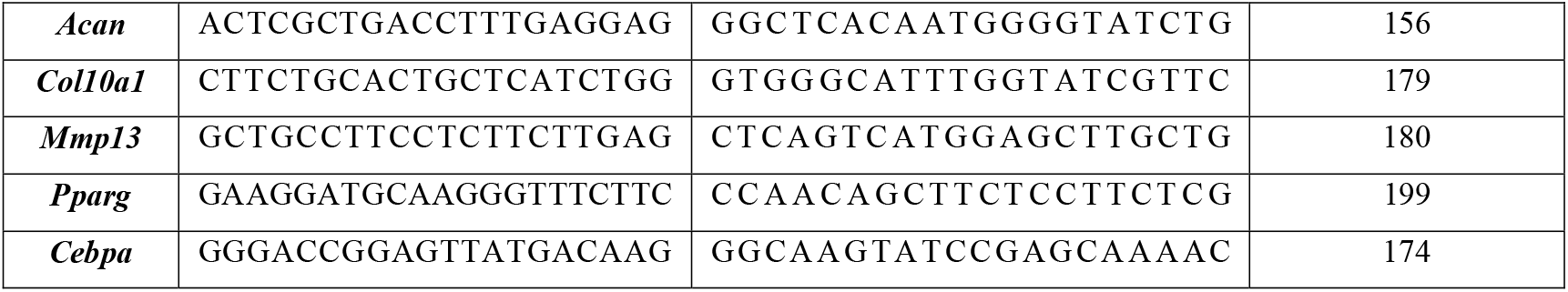
Primer List.

### Staining

The spheroids were collected and fixed in 4% paraformaldehyde (PFA). After fixation, the samples were permeabilized with 0.1% triton X-100 in PBS for 30 min and washed with PBS 3 times. For immunostaining, non-specific binding during staining was blocked with PBS containing 4% bovine serum albumin at room temperature for 1 h. Then, we added the anti-SOST antibody (Sigma) and incubated at room temperature for 1 h. After washing with PBS 3 times, the samples were treated with the Alexa Flour 488 secondary antibody (Invitrogen), Alexa Fluor 546 Phalloidin (Invitrogen), and DAPI (Sigma) at room temperature for 1 h. After washing with PBS 3 times, the samples were observed by the FLUOVIEW FV3000 (Olympus). For G-actin (globular) and F-actin (fibrous) staining, the samples were treated with the Deoxyribonuclease I (DNase I), Alexa Fluor 488 conjugate (Invitrogen) and Alexa Fluor 546 Phalloidin (Invitrogen), respectively, and counterstained with DAPI (Sigma) at room temperature for 15 min. After washing with PBS 3 times, the stained samples were mounted on the glass slides for visualization.

### Statistical analysis

The statistical significance was assessed by Student’s t-test, F-test, or ANOVA followed by Tukey’s honestly significant difference (HSD) *post-hoc* test (with α = 0.05). P-values less than 0.05 were considered to be significant.

## Result

### MSC spheroid with osteogenic induction supplements up-regulated osteocyte markers within 2 days

As described in Fig. 1(A), the spheroids reconstructed by hMSCs were incubated in the osteogenic induction (OI) medium for 2 days. In Fig. 1(B), the spheroid has about 185 μm diameter at 2-day. To evaluate gene expression changes in the spheroid compared to the monolayer, we carried out real-time PCR in Fig. 1(C). After 2-day incubation, the relative mRNA expression in the spheroid for stem cell markers were up-regulated compared to that in the monolayer; *Nanog* (22.6-fold change; *p* < 0.05), *Oct4* (9.7-fold change; *p* < 0.005), and *Sox2* (30.4-fold change; *p* < 0.05). Regarding osteoblast markers, there was no significant mRNA expression change in *Runx2* (1.70-fold change), while both *Alpl* (0.41-fold change; *p* < 0.005) and *Col1a1* (0.21-fold change; *p* < 0.005) mRNA expressions were down-regulated in the spheroid. On the other hand, the relative mRNA expressions for osteocyte markers in the spheroids compared to the monolayer were highly up-regulated; *Opn* (28.4-fold change; *p* < 0.05), *Phex* (18.4-fold change; *p* < 0.005), and *Sost* (21.7-fold change; *p* < 0.05). To examine the protein level expression of osteocyte marker, we conducted the SOST immunostaining in Fig. 1(D) since the SOST is known as a marker of mature osteocytes. As a result, the SOST expression (green) in the protein level was detected entirely inside the spheroid after 2-day incubation. Moreover, F-actin was expressed strongly on the surface of the spheroid compared to the inner part.

**Figure 1.**
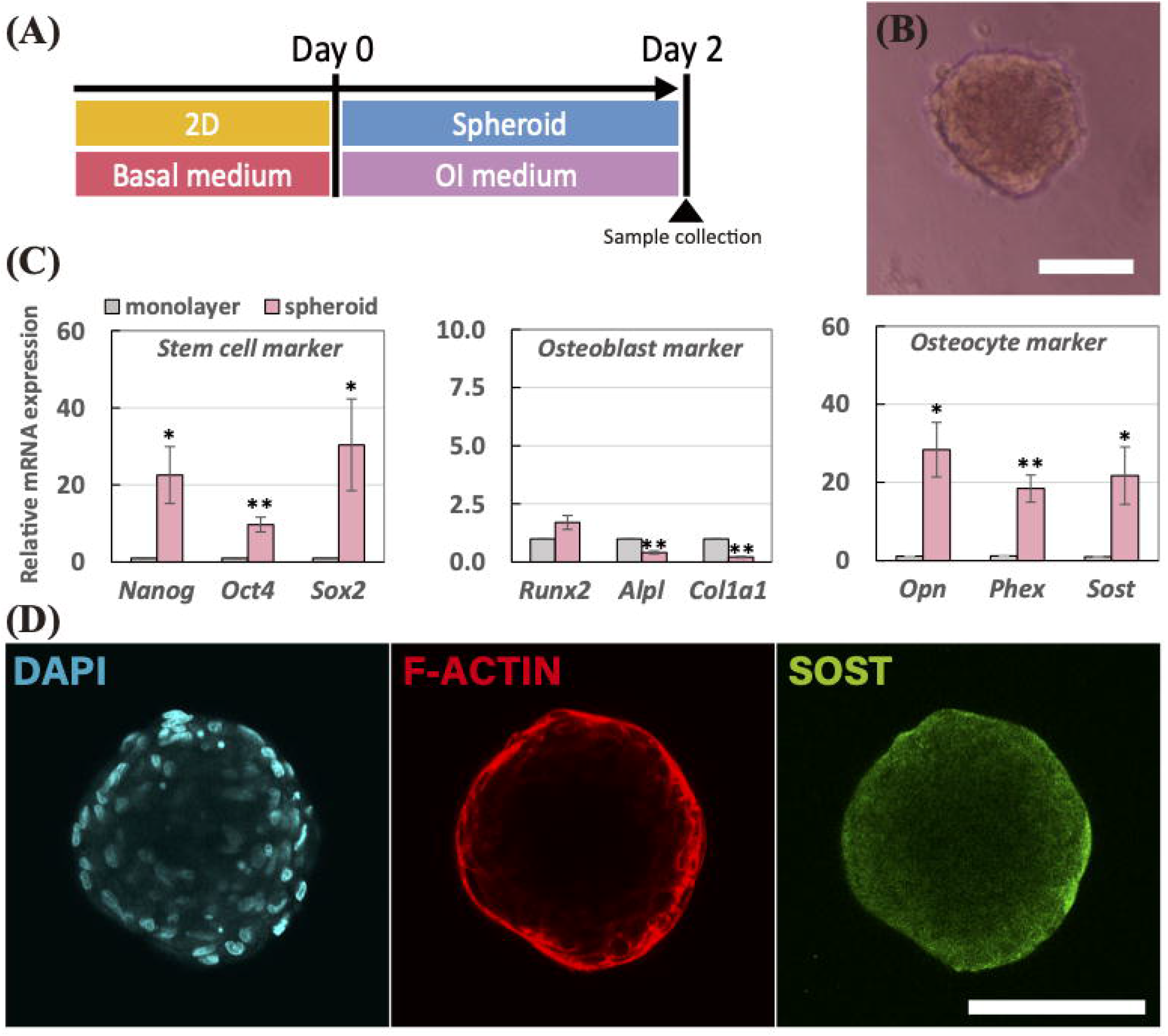
(A) Schematic of the experimental time line for spheroids incubated in the osteogenesis induction medium for 2 days. (B) Morphology of the spheroid after a 2-day incubation period. The scale bar represents 100 μm. (C) Relative mRNA expressions of stem cell markers (*Nanog*, *Oct4*, and *Sox2*), osteoblast markers (*Runx2*, *Alpl*, and *Col1a1*), and osteocyte markers (*Opn*, *Phex*, and *Sost*) were measured by real-time PCR. All the mRNA expressions were normalized to *Gapdh* expressions while the results were expressed as relative amounts against the expression of monolayer sample (*n* = 8). The bars represent the mean ± standard error. *P*-value was calculated from Student’s *t*-test; \**p* < 0.05, \*\**p* < 0.005. (D) Immunostaining images of spheroid after 2-day incubation; cell nuclei (DAPI in cyan), actin filaments (F-Actin in red), and sclerostin (SOST in green). The scale bar represents 100 μm.

### Up-regulated osteocyte markers in the spheroid were prolonged after 7 days

To investigate the long-term persistence of gene expressions for osteocyte markers in the spheroid, we performed the longer experiment, 7-day, as shown in Fig.2 (A). The spheroid at 7-day in Fig. 2(B) became smaller compared to 2-day spheroid (48.9% reduction; *p* < 0.005). In Fig. 2 (C), the mean values of projected area of spheroid at 2-day and 7-day were 25,900 μm^2^ and 12,700 μm^2^, respectively. In Fig. 2(D), we conducted real-time PCR to examine the gene expression changes between 2-day and 7-day spheroid. Consequently, the stem cell markers in the 7-day spheroid were up-regulated compared to the 2-day spheroid; *Nanog* (15.9-fold change; *p* = 0.07), *Oct4* (12.8-fold change; *p* < 0.05), and *Sox2* (12.7-fold change; *p* = 0.26). Osteoblast markers were non-significantly modulated; *Runx2* (2.63-fold change; *p* = 0.18), *Alpl* (8.44-fold change; *p* = 0.09), and *Col1a1* (0.68-fold change; *p* = 0.12). Regarding osteocyte markers, while early osteocyte marker, *Opn* (0.86-fold change; *p* = 0.75), was not modulated, it up-regulated the mature osteocyte markers in the 7-day spheroid including *Phex* (18.5-fold change; *p* < 0.05) and *Sost* (10.0-fold change; *p* < 0.005). Immunostaining results in Fig. 2(E) represented that the SOST expression was entirely expressed in the 7-day spheroid

**Figure 2.**
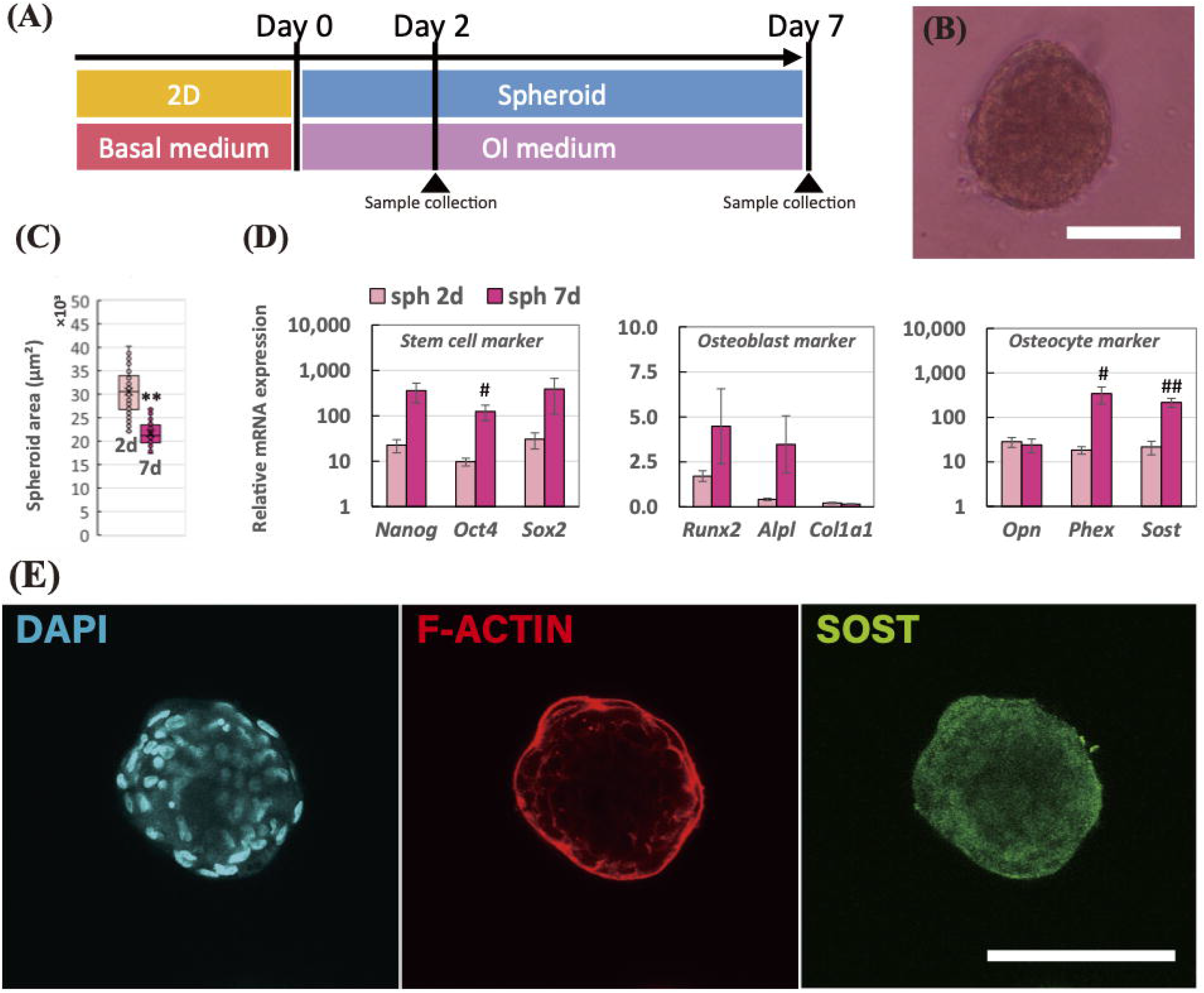
(A) Schematic of the experimental time line for spheroids incubated in the osteogenesis induction medium for 7 days. (B) Morphology of the spheroid after a 7-day incubation period. The scale bar represents 100 μm. (C) Mean values of projected area of the spheroids after 2 and 7 days. F-test was performed to examine the significance of area changes in spheroids (*n* = 60 from 5 independent experiments; \**p* < 0.05, \*\**p* < 0.005). (D) Relative mRNA expressions of stem cell markers (*Nanog*, *Oct4*, and *Sox2*), osteoblast markers (*Runx2*, *Alpl*, and *Col1a1*), and osteocyte markers (*Opn*, *Phex*, and *Sost*) in the spheroids at 2-day and 7-day incubation were measured by real-time PCR. All the mRNA expressions were normalized to *Gapdh* expressions while the results were expressed as relative amounts against the expression of monolayer sample (*n* = 8). The bars represent the mean ± standard error. *P*-value was calculated from Student’s *t*-test; ^#^*p* < 0.05, ^##^*p* < 0.005. (E) Immunostaining images of spheroid after 2-day incubation; cell nuclei (DAPI in cyan), actin filaments (F-actin in red), and sclerostin (SOST in green). The scale bar represents 100 μm.

### Structural effect achieved in 3D spheroid compared to 2D monolayer promoted stem cell, osteocyte, and hypertrophic chondrocyte markers

As described in Fig. 3(A), the spheroids were incubated in the basal medium in the absence of OI medium in order to elucidate the structural effect achieved in the 3D spheroid culture compared to the 2D monolayer condition, without the chemical supplements. In Fig. 3(B), we measured the gene expression changes in stem cell, osteoblast, and osteocyte markers. As a result, all the stem cell markers were significantly up-regulated in both 2-day and 7-day spheroid compared to 2-day monolayer, corresponding to the previous experiments. Particularly, the stem cell markers in the spheroid were further up-regulated after 7-day incubation. Regarding the osteoblast markers, there was significant up-regulations in the monolayer model, whereas those markers in the spheroid exerted a different trend. Especially, the *Col1a1* mRNA expressions were significantly down-regulated at both 2-day and 7-day. In the spheroid model, all the osteocyte markers were up-regulated at 2-day and 7-day. Particularly, mRNA expression of early osteocyte marker, *Opn,* in 2-day spheroid was higher than that in 7-day spheroid, whereas the mRNA expressions of matured osteocyte markers involving *Phex* and *Sost* in 7-day spheroid was greater than that in 2-day spheroid.

**Figure 3.**
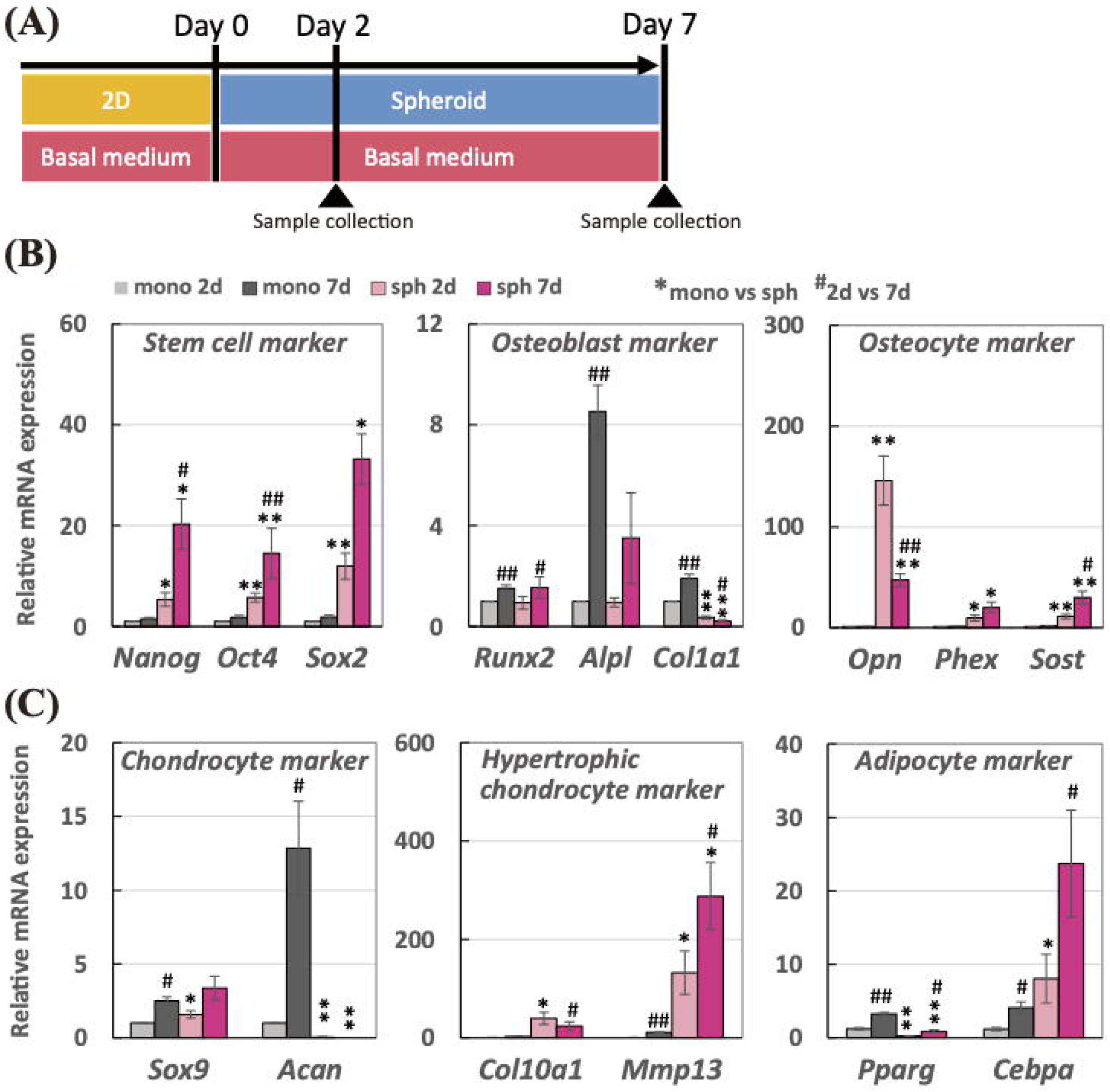
(A) Schematic of the experimental time line for spheroids incubated in the basal medium in the absence of osteogenesis supplements for 7 days. Relative mRNA expressions of (B) stem cell markers (*Nanog*, *Oct4*, and *Sox2*), osteoblast markers (*Runx2*, *Alpl*, and *Col1a1*), osteocyte markers (*Opn*, *Phex*, and *Sost*), (C) chondrocyte markers (*Sox9* and *Acan*), hypertrophic chondrocyte marker (*Col10a1* and *Mmp13*), and adipocyte markers (*Pparg* and *Cebpa*) were measured by real-time PCR. All the mRNA expressions were normalized to *Gapdh* expressions while the results were expressed as relative amounts against the expression of monolayer sample (*n* = 10). The bars represent the mean ± standard error. *P*-value was calculated from Student’s *t*-test (mono vs sph: \**p* < 0.05, \*\**p* < 0.005; 2-day vs 7-day: ^#^*p* < 0.05, ^##^*p* < 0.005).

Since the differentiation of hMSCs was not chemically induced into a certain cell lineage in this experiment, we also examined the gene expression change in chondrocyte, hypertrophic chondrocyte, and adipocyte markers. In Fig. 3(C), there was no significant changes in the osteocyte gene expression in the monolayer model. Regarding chondrocyte markers, both *Sox9* and *Acan* mRNA expressions were up-regulated in the monolayer condition after 7-day incubation. In the spheroid, however, *Acan* mRNA expression was greatly suppressed at both 2-day and 7-day incubation although *Sox9* mRNA expression was slightly increased. On the other hand, hypertrophic chondrocyte markers, such as *Col10a1* and *Mmp13*, were greatly up-regulated in the form of spheroid compared to the monolayer. For adipose markers, the adipogenesis master gene, *Pparg*, was greatly down-regulated in the spheroid although mRNA expression of the metabolic action of insulin marker, *Cebpa,* was up-regulated in the spheroid. On the other hand, both *Pparg* and *Cebpa* mRNA expressions were increased in the monolayer condition after 7-day incubation.

### Cell condensed condition in the 3D spheroid structure sustained the osteocyte differentiation

To elucidate the effect of cell condensation achieved in the 3D spheroid structure on the osteocyte differentiation, we introduced another model which the 3D spheroids in the ultra-low attachment dish at 2-day incubation were transferred to a normal culture dish as described in Fig. 4(A). Immediate after the spheroid was subcultured on the normal culture dish, the spheroids initiated to attach on the dish in Fig 4(B), and then the cells dissociated from the spheroid spread over the dish. After the extra 2-day incubation on the normal culture dish, the spheroid at 2-day lost its 3D structure and became like 2D model in Fig. 4(C), termed sph-mono model. We then examined their gene expression levels in the sph-mono model compared to 4-day monolayer and spheroid using real-time PCR in Fig. 4(D). Similar to 2-day cultivation results in the previous experiment, 4-day spheroid had higher gene expression levels in the stem cell and osteocyte markers than 4-day monolayer, whereas the mRNA expressions of osteoblast markers such as *Alpl a*nd *Col1a1* in the spheroid were suppressed compared to the monolayer. On the other hand, those up-regulated markers for stem cell and osteocytes were down-regulated in the sph-mono model, which indicate that the up-regulations of stem cell and osteocyte markers were depending on the cell condensed condition acquired from the 3D spheroid structure. Regarding osteoblast markers, there was no significant change in the sph-mono model compared to the spheroid.

**Figure 4.**
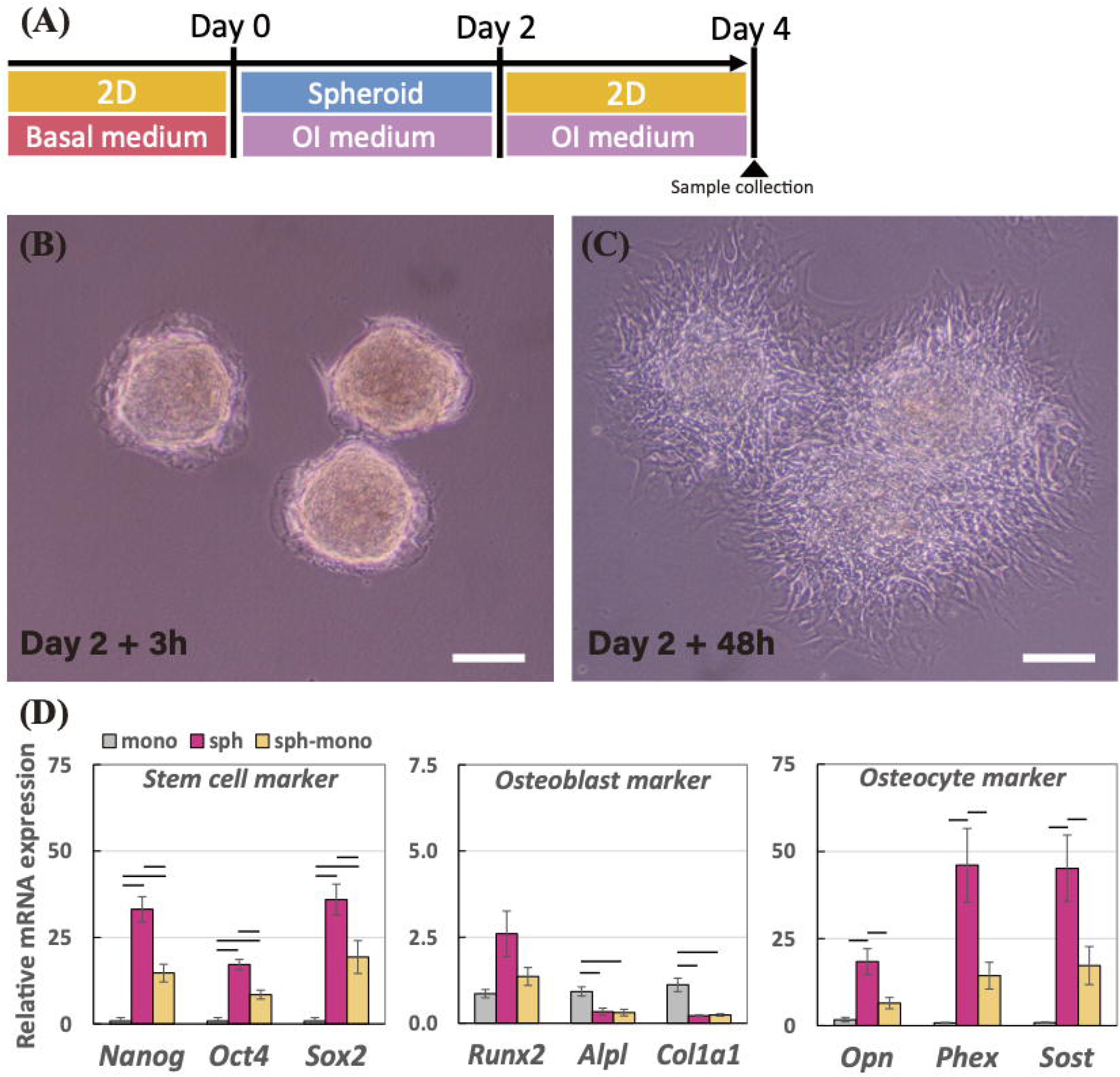
(A) Schematic of the experimental time line to prepare for cells dissociated from the spheroids (sph-mono) in the osteogenesis induction medium. Morphology of the spheroids transferred to the normal culture dish after (B) 3 h and (C) 48 h. The scale bar represents 200 μm. (D) Relative mRNA expressions of stem cell markers (*Nanog*, *Oct4*, and *Sox2*), osteoblast markers (*Runx2*, *Alpl*, and *Col1a1*), and osteocyte markers (*Opn*, *Phex*, and *Sost*) in monolayer and spheroid incubated for 4 days, and sph-mono were measured by real-time PCR. All the mRNA expressions were normalized to *Gapdh* expressions while the results were expressed as relative amounts against the expression of monolayer sample (*n* = 6). The graphs represent the mean ± standard error. Bar indicates the significance between groups, which was calculated from ANOVA with Tukey’s HSD *post-hoc* test (α = 0.05).

### Cytochalasin D facilitated the osteocyte differentiation in the spheroid by actin depolymerization

To reveal mechanisms for the osteocyte differentiation evoked in the 3D spheroid structure, we utilized an actin polymerization inhibitor, cytochalasin D. During fabrication of spheroid, we added 1 μM of cytochalasin D as described in Fig. 5(A). Compared to the spheroid with DMSO in Fig. 5(B), the size of spheroid treated with cytochalasin D became bigger as shown in Fig. 5(C). In Fig. 5(D), the mean values of projected area of spheroid in the presence of DMSO and cytochalasin D became 23,500 μm^2^ and 31,300 μm^2^, respectively (133% increase; *p* < 0.005). In Fig. 5(E), we carried out F-actin/G-actin staining using phalloidin and DNase I to examine the changes in the actin balancing. By addition of cytochalasin D in the spheroid, the F-actin represented among the cells in the spheroid was delocalized. Moreover, the spheroid treated with cytochalasin D exerted greater G-actin expressions compared to the spheroid with DMSO. In Fig. 5(F), we then conducted real-time PCR to examine the mRNA expression changes in the spheroid by addition of inhibitor. Among the stem cell markers, adding cytochalasin D significantly up-regulated *Sox2* mRNA expression; *Nanog* (1.15-fold change), *Oct4* (1.13-fold change), and *Sox2* (1.68-fold change; *p* < 0.05). Moreover, osteoblast markers were modulated by cytochalasin D; *Runx2* (1.40-fold change; *p* < 0.005), *Alpl* (2.95-fold change; *p* < 0.05), and *Col1a1* (0.82-fold change; *p* < 0.05). Particularly, addition of cytochalasin D up-regulated the osteocyte mRNA expressions; *Opn* (3.80-fold change; *p* < 0.05), *Phex* (1.15-fold change), *Sost* (1.24-fold change; *p* < 0.05). Hence, actin depolymerization by cytochalasin D in the spheroid up-regulated the osteocyte gene expression levels in the spheroid.

**Figure 5.**
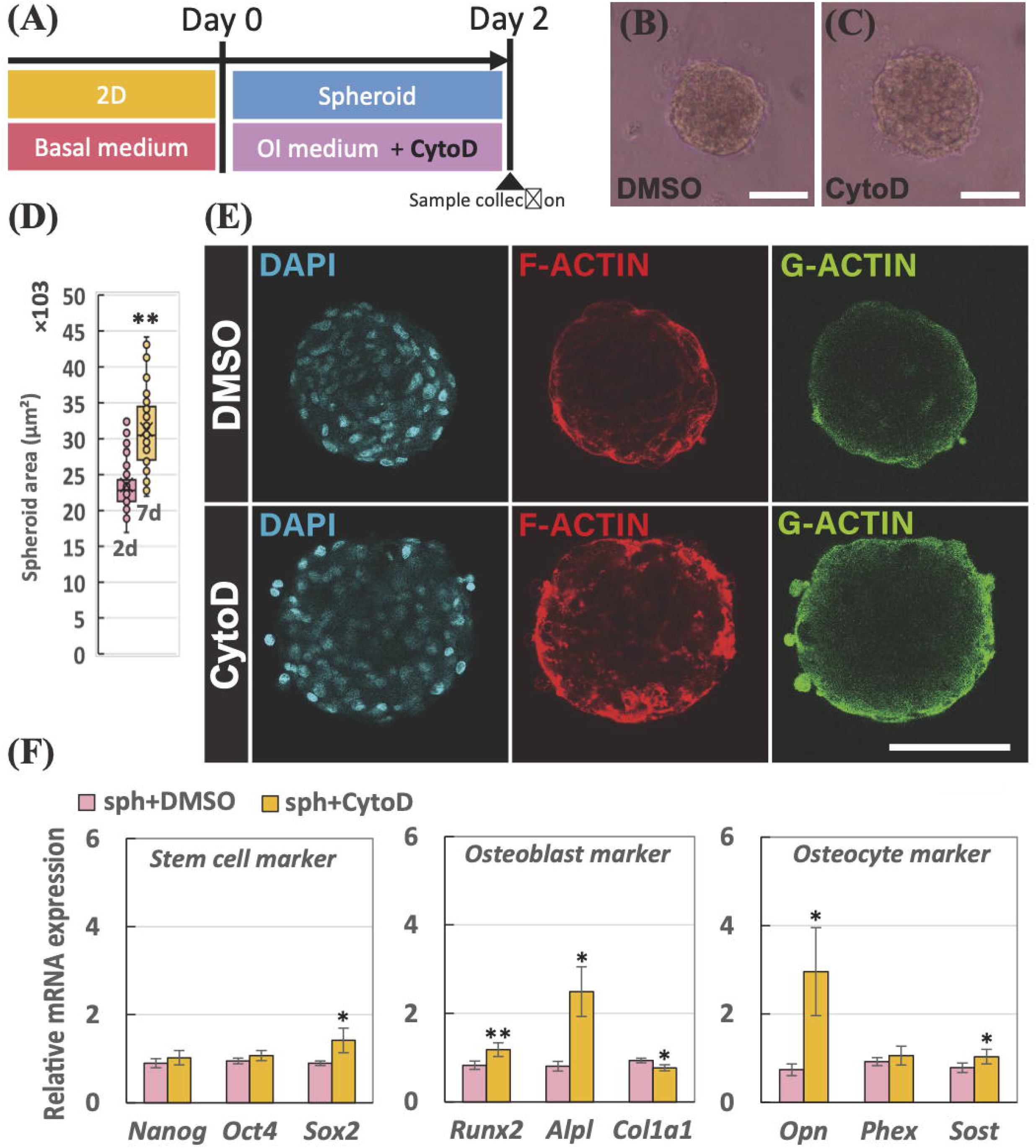
(A) Schematic of the experimental time line to fabricate the spheroid treated with 1 μM of cytochalasin D (CytoD). Morphology of the spheroids treated with (B) DMSO and (C) CytoD. The scale bar represents 100 μm. (D) Mean values of projected area of the spheroids treated with DMSO and CytoD incubated for 2 days. (E) Staining images of spheroid after 2-day incubation; cell nuclei (DAPI in cyan), fibrous actin (F-actin in red), and globular actin (G-actin in green). The scale bar represents 100 μm. (F) Relative mRNA expressions of stem cell markers (*Nanog*, *Oct4*, and *Sox2*), osteoblast markers (*Runx2*, *Alpl*, and *Col1a1*), and osteocyte markers (*Opn*, *Phex*, and *Sost*) in the spheroid treated with DMSO and CytoD were measured by real-time PCR. All the mRNA expressions were normalized to *Gapdh* expressions while the results were expressed as relative amounts against the expression of spheroid samples treated with DMSO (*n* = 9). The bars represent the mean ± standard error. *P*-value was calculated from Student’s *t*-test; \**p* < 0.05, \*\**p* < 0.005.

## Discussion

Our group has studied the osteocyte differentiation acquired from the 3D culture system. In the previous studies, we showed that the osteocyte differentiation of pre-osteoblast cells was facilitated in the 3D scaffold-free culture system (Kim and Adachi, 2019; Kim et al., 2020; Kim and Adachi, 2020). Although we revealed that cell condensed condition is significant for the pre-osteoblast cells to undergo the osteocyte differentiation, it is still unclear whether or not the osteocyte differentiation can be further evoked from the mesenchymal stem cells that recapitulate the *in vitro* bone development process. In this study, we assumed that the 3D spheroid culture reconstructed by hMSCs enables to recapitulate the mesenchymal condensation process during the ossification. By using hMSCs, we fabricated self-organized spheroids and evaluated its osteocyte-likeness *in vitro*.

Under the OI medium, the hMSC spheroids were chemically induced to differentiate into the osteogenic lineage. As a result of real-time PCR and SOST immunostaining in Figs 1 and 2, the cells in the spheroid successfully rendered osteocyte-likeness within 2 days while the osteocyte-likeness was further prolonged up to 7 days. Furthermore, the osteoblast markers were greatly suppressed in the spheroid compared to the monolayer condition. The results obtained with the hMSCs in the present study corresponded to the previous studies using the murine pre-osteoblast cells, MC3T3-E1 (Kim and Adachi, 2019; Kim and Adachi, 2020). Moreover, the spheroid also exerted the up-regulations in stem cell markers. The increase in stem cell markers in the spheroid culture were also reported from other studies using stem cells (Cheng et al., 2012; Guo et al., 2014; Zhou et al., 2017). While the osteoblast differentiation in the monolayer condition is actively proceeded, the stemness were relatively maintained in the spheroid culture, so that the stem cell gene expressions were relatively up-regulated in the spheroid culture.

We then conducted the experiment with the basal medium for 2 and 7 days in the absence of OI supplements in order to investigate the independent structural effect in the 3D spheroid culture against the 2D monolayer culture. Interestingly, the differentiation induction of hMSCs had distinguished trends depending on the cultivation condition. In the monolayer condition, the hMSCs were likely to undertake the osteoblast, chondrocyte, and adipocyte differentiation randomly because of the absence of specific differentiation induction mediums. The result suggested that the monolayer condition seems to be a suitable condition to conduct the osteoblast, chondrocyte, and adipocyte differentiation studies as conducted in other studies for several decades. Since no certain chemical supplements were utilized, the hMSCs were thought to differentiate into those cells randomly. While the differentiation processes into osteoblast, chondrocyte, and adipocyte were undertaken on the monolayer, the spheroid condition had relatively greater gene expressions in osteocyte and hypertrophic chondrocyte markers as well as stem cell markers. During the endochondral ossification process, it is known that the bone is replaced from the cartilage via hypertrophic chondrocytes (Aghajanian and Mohan, 2018). The transient up-regulations of osteocyte and hypertrophic chondrocyte markers in the spheroid model might mimic the certain event of ossification process *in vitro* which is initiated from the mesenchymal condensation *in vivo* during the bone development process. Regarding the stem cell markers, the increase in stem cell gene expressions were observed, corresponding to the previous experiment using the OI medium in Fig. 1, which indicates that the spheroid culture for the hMSC relatively maintained its stemness compared to the monolayer condition, whereas the cells in the monolayer model is likely to differentiate and commit to some lineages even without certain chemical supplements.

To investigate the cell condensed effect achieved by the 3D spheroid, we transferred the spheroid to the normal culture dish, causing that the 3D spheroid spread over and became like monolayer condition, termed sph-mono model (the cells dissociated from the 3D spheroid). In terms of gene expression changes in the sph-mono model compared to 4-day spheroid and monolayer, the up-regulated osteocyte mRNA expressions as well as stemness marker in the spheroids were diminished after the spheroid lost the condensed condition. The results suggested that the cell condensed condition is essential for the hMSCs to differentiate into osteocytes and further to retain its osteocyte-likeness. In other words, the monolayer condition is not a suitable cell cultural environment for osteocyte related studies. This is why only limited number of *in vitro* osteocytic model were available since many studies attempted to undertake osteocyte differentiation on the monolayer condition. Hence, we hereby highlighted the significance of cell condensation condition for osteocytogenesis.

In order to reveal the mechanism in the facilitated osteocyte differentiation in the spheroid structure, the involvement of actin balancing was examined. In this study, cytochalasin D was utilized to inhibit the actin polymerization in the spheroid. As a result, cytochalasin D successfully delocalized F-actin in the spheroid, resulting in the increase in the size of spheroid. The prevention of F-actin polymerization by cytochalasin D eventually promoted osteocyte mRNA expressions in the spheroid. With regard to osteoblast differentiation on the conventional 2D monolayer condition, the actin balancing is involved in the osteoblast differentiation of human bone marrow-derived MSCs (Chen et al., 2015). On the other hand, our results first highlighted the involvement of actin balancing for the osteocyte differentiation of hMSCs. As illustrated in Fig. 6, this study suggested that the osteocyte differentiation can be achieved by weakened F-actin formation in the spheroid compared to the monolayer, whereas the generation of the tight F-actin on the monolayer model evoked the osteoblast differentiation of hMSCs as well as bone mineralization as shown in many other studies. For the stemness markers, actin depolymerization in the stem cells is known to promote their pluripotent states (Yu et al., 2018), which corresponds to our results. Based on this inhibitor experiment using the spheroid model, we first showed that the actin balancing is involved in the osteocyte differentiation induction acquired by the spheroid culture.

**Figure 6.**
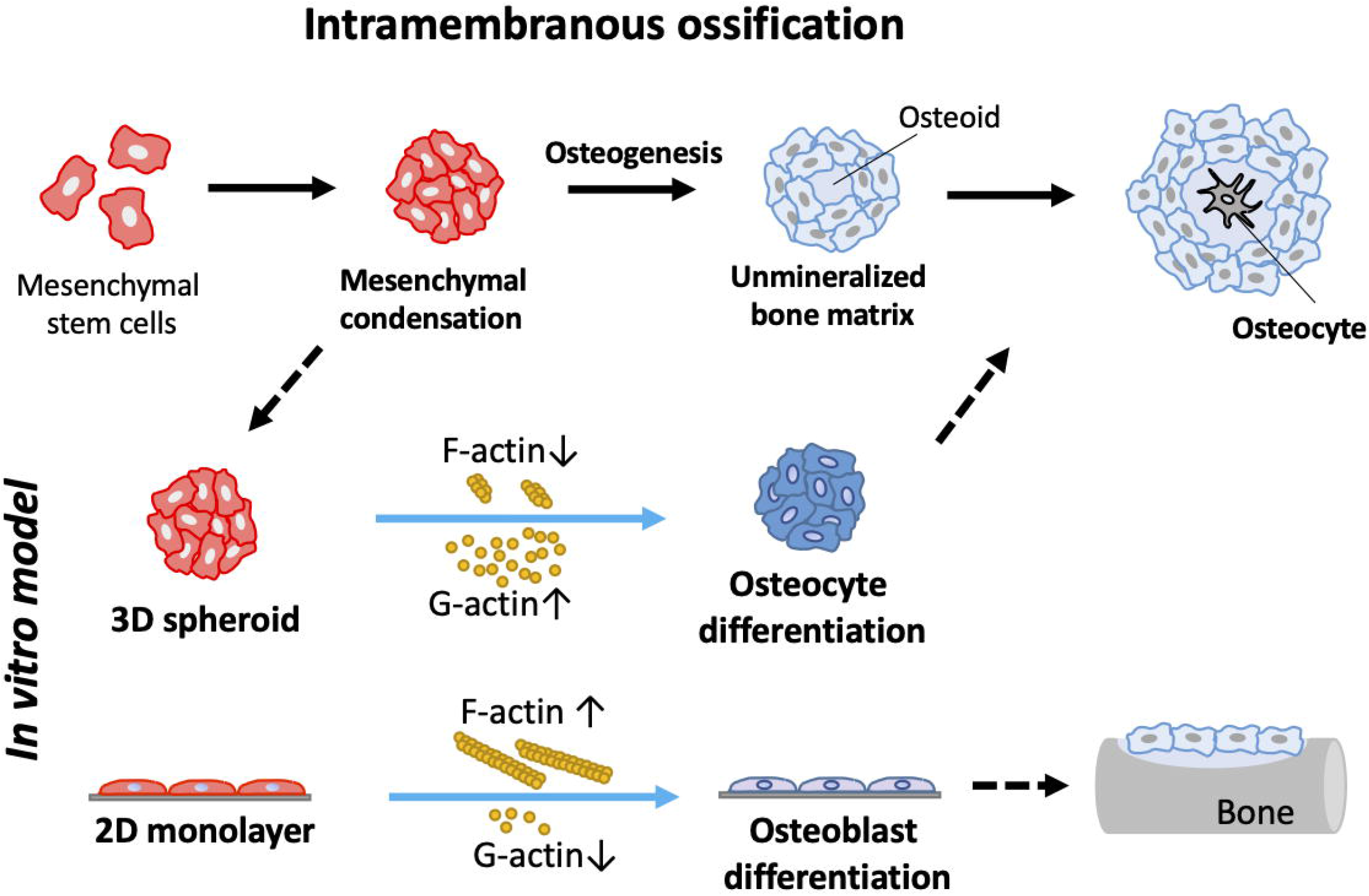
Schematic diagram to highlight the significance of actin balancing in the 3D spheroid culture to determine the cell differentiation fate into osteocytes.

As many studies reported, the osteoblasts undergo the bone mineralization process where they are two-dimensionally arranged on the bone matrix, which is reconstructed in the conventional 2D monolayer condition for a long period of cultivation time, from several weeks to several months. During the *in vitro* bone formation studies using the 2D model, the osteoblast precursor cells in the confluent monolayer condition eventually forms the 3D dome-shape of bone nodules after a long experiment over several weeks (Bhargava et al., 1988). Eventually, osteocyte-like cells were found in the mineralized nodule (Kawai et al., 2019). Throughout our present study using the 3D spheroid culture model, we suggested a new insight that the osteocyte differentiation can be triggered by the 3D cell condensed condition of osteoblast precursor cells within a short period of time at a certain remodeling site. It might indicate that a group of cells in the bone nodule are subjected to the cell condensed condition, so that the osteocyte differentiation is acquired after the osteoblast precursor cells are confronted with the cell condensed condition. This explanation might also imply that the osteoblasts in the bone become osteocytes when they encountered the condensed condition at the remodeling site *in vivo*. For the cells in the condensed condition, they became three-dimensionally surrounded by cells and their extracellular matrices, so that formation of actin filaments is thought to be greatly suppressed. Hence, the osteocyte might not be differentiated from the mature osteoblasts, but directly differentiated from the osteoblast precursor cells in the cell condensed condition. Consequently, the cell condensed condition plays a significant role to determine the cellular differentiation fate of MSCs into the osteocytes.

In this study, the 3D spheroid reconstructed by MSCs under the OI medium exerted the osteocyte-likeness within 2 days. Notably, we first suggested the significance of the environmental factor to determine the MSC differentiation fate to osteocytes or osteoblasts, such as cell condensed condition acquired from the 3D culture beyond the chemically induced supplements. Moreover, we investigated the structural effect on MSC differentiations, which provides a suitable strategy of the cultural model for certain cell lineage studies. In the future, our 3D spheroid culture model can be utilized to represent the osteocytes and further to recapitulate an *in vitro* ossification process, becoming a new *in vitro* model as “bone organoid”.

## Acknowledgement

We would like to thank Junko Sunaga for providing technical supports. This work was supported by the Japan Society for the Promotion of Science (JSPS) KAKENHI (20H00659, 20K20181, and 19K23604), Advanced Research and Development Programs for Medical Innovation (AMED-CREST), by elucidation of mechanobiological mechanisms and their application to the development of innovative medical instruments and technologies from Japan Agency for Medical Research and Development (AMED) (JP20gm0810003), and by the acceleration program for intractable diseases research utilizing disease-specific iPS cells from AMED (JP19bm0804006).

## References

1. Adachi T, Aonuma Y, Ito S, Tanaka M, Hojo M, Takano-Yamamoto T, Kamioka H. 2009. Osteocyte calcium signaling response to bone matrix deformation. J Biomech 42:2507–2512. doi:10.1016/j.jbiomech.2009.07.006

2. Aghajanian P, Mohan S. 2018. The art of building bone: Emerging role of chondrocyte-to-osteoblast transdifferentiation in endochondral ossification. Bone Res 6. doi:10.1038/s41413-018-0021-z

3. Bhargava U, Bar-Lev M, Bellows CG, Aubin JE. 1988. Ultrastructural analysis of bone nodules formed in vitro by isolated fetal rat calvaria cells. Bone 9:155–163. doi:10.1016/8756-3282(88)90005-1

4. Bonewald LF. 2006. Mechanosensation and transduction in osteocytes. BoneKEy-Osteovision 3:7–15. doi:10.1138/20060233

5. Buttery LDK, Bourne S, Xynos JD, Wood H, Hughes FJ, Hughes SPF, Episkopou V, Polak JM. 2001. Differentiation of osteoblasts and in Vitro bone formation from murine embryonic stem cells. Tissue Eng. 7:89–99. doi:10.1089/107632700300003323

6. Caplan AI. 1991. Mesenchymal stem cells. J Orthop Res. 9(5):641−650. doi:10.1002/jor.1100090504

7. Chen L, Shi K, Frary CE, Ditzel N, Hu H, Qiu W, Kassem M. 2015. Inhibiting actin depolymerization enhances osteoblast differentiation and bone formation in human stromal stem cells. Stem Cell Res 15:281–289. doi:10.1016/j.scr.2015.06.009

8. Cheng NC, Wang S, Young TH. 2012. The influence of spheroid formation of human adipose-derived stem cells on chitosan films on stemness and differentiation capabilities. Biomaterials 33:1748–1758. doi:10.1016/j.biomaterials.2011.11.049

9. Coelho MJ, Fernandes MH. 2000. Human bone cell cultures in biocompatibility testing. Part II: Effect of ascorbic acid, β-glycerophosphate and dexamethasone on osteoblastic differentiation. Biomaterials 21:1095–1102. doi:10.1016/S0142-9612(99)00192-1

10. Guo L, Zhou Y, Wang S, Wu Y. 2014. Epigenetic changes of mesenchymal stem cells in three-dimensional (3D) spheroids. J Cell Mol Med 18:2009–2019. doi:10.1111/jcmm.12336

11. Kawai S, Yoshitomi H, Sunaga J, Alev C, Nagata S, Nishio M, Hada M, Koyama Y, Uemura M, Sekiguchi K, Maekawa H, Ikeya M, Tamaki S, Jin Y, Harada Y, Fukiage K, Adachi T, Matsuda S, Toguchida J. 2019. In vitro bone-like nodules generated from patient-derived iPSCs recapitulate pathological bone phenotypes. 3: 558–570. Nat Biomed Eng. doi:10.1038/s41551-019-0410-7

12. Kim J, Adachi T. 2019. Cell Condensation Triggers the Differentiation of Osteoblast Precursor Cells to Osteocyte-Like Cells. Front Bioeng Biotechnol 7:288. doi:10.3389/fbioe.2019.00288

13. Kim J, Kigami H, Adachi T. 2020. Characterization of self-organized osteocytic spheroids using mouse osteoblast-like cells. J Biomech Sci Eng 15:1–8. doi:10.1299/jbse.20-00227

14. Kim J, Adachi T. 2020. Modulation of Sost gene epxression under hypoxia in 3D scaffold-free osteocytic tissue. Tissue Eng Part A In press. doi:10.1089/ten.tea/2020.0228

15. Manolagas SC. 2000. Birth and death of bone cells: Basic regulatory mechanisms and implications for the pathogenesis and treatment of osteoporosis. Endocr Rev 21:115–137. doi:10.1210/er.21.2.115

16. Mechiche Alami S, Gangloff SC, Laurent-Maquin D, Wang Y, Kerdjoudj H. 2016. Concise Review: In Vitro Formation of Bone - Like Nodules Sheds Light on the Application of Stem Cells for Bone Regeneration. Stem Cells Transl Med. 5(11):1587–1593. doi:10.5966/sctm.2015-0413

17. Parfitt AM. 2001. The bone remodeling compartment: A circulatory function for bone lining cells. J Bone Miner Res 16:1583–1585. doi:10.1359/jbmr.2001.16.9.1583

18. Pittenger MF, Discher DE, Péault BM, Phinney DG, Hare JM, Caplan AI. 2019. Mesenchymal stem cell perspective: cell biology to clinical progress. npj Regen Med 4:22. doi:10.1038/s41536-019-0083-6

19. Pittenger MF, Mackay AM, Beck SC, Jaiswal RK, Douglas R, Mosca JD, Moorman MA, Simonetti DW, Craig S, Marshak DR. 1999. Multilineage potential of adult human mesenchymal stem cells. Science 284(5411):143–147. doi:10.1126/science.284.5411.143

20. Robling AG, Bonewald LF. 2020. The Osteocyte□: New Insights. Annu Rev Physiol 82:485–506. doi:10.1146/annurev-physiol-021119-034332

21. Rutkovskiy A, Stensløkken K-O, Vaage IJ. 2016. Osteoblast Differentiation at a Glance. Med Sci Monit Basic Res 22:95–106. doi:10.12659/msmbr.901142

22. Yu L, Li J, Hong J, Takashima Y, Fujimoto N, Nakajima M, Yamamoto A, Dong X, Dang Y, Hou Y, Yang W, Minami I, Okita K, Tanaka M, Luo C, Tang F, Chen Y, Tang C, Kotera H, Liu L. 2018. Low Cell-Matrix Adhesion Reveals Two Subtypes of Human Pluripotent Stem Cells. Stem Cell Reports. 11(1): 142–156. doi:10.1016/j.stemcr.2018.06.003

23. Zhou Y, Chen H, Li H, Wu Y. 2017. 3D culture increases pluripotent gene expression in mesenchymal stem cells through relaxation of cytoskeleton tension. J Cell Mol Med 21:1073–1084. doi:10.1111/jcmm.12946

